# Structural basis for σ_1_ receptor ligand recognition

**DOI:** 10.1101/333765

**Authors:** Hayden R. Schmidt, Robin M. Betz, Ron O. Dror, Andrew C. Kruse

## Abstract

The σ_1_ receptor is a poorly understood integral membrane protein expressed in most cells and tissues in the human body. It has been shown to modulate the activity of other membrane proteins such as ion channels and G protein-coupled receptors^1–4^, and ligands targeting the σ_1_ receptor are currently in clinical trials for treatment of Alzheimer’s disease^5^, ischemic stroke^6^, and neuropathic pain^7^. Despite its importance, relatively little is known regarding σ_1_ receptor function at the molecular level. Here, we present crystal structures of the human σ_1_ receptor bound to the classical antagonists haloperidol and NE-100, as well as the agonist (+)-pentazocine, at crystallographic resolutions of 3.1 Å, 2.9 Å, and 3.1 Å respectively. These structures reveal a unique binding pose for the agonist. The structures and accompanying molecular dynamics (MD) simulations demonstrate that the agonist induces subtle structural rearrangements in the receptor. In addition, we show that ligand binding and dissociation from σ_1_ is a multistep process, with extraordinarily slow kinetics limited by receptor conformational change. We use MD simulations to reconstruct a ligand binding pathway that requires two major conformational changes. Taken together, these data provide a framework for understanding the molecular basis for agonist action at σ_1_.

---

Discovered in 1976^8^, the σ_1_ receptor has attracted interest because it binds a host of structurally dissimilar pharmacologically active compounds with high affinity (Fig. 1a). These include benzomorphans, antipsychotics, psychosis-inducing drugs, the antifungal agent fenpropimorph, sterols such as progesterone, and numerous other compounds^9^. These molecules contain few shared features, although most include a basic nitrogen atom flanked on two sides by longer hydrophobic moieties (typically phenyl rings), representing a minimal σ_1_-binding pharmacophore (Fig. 1a)^10^. Cloning of the σ_1_ receptor showed that it bears no similarity to any other human protein^11^. Instead, its nearest homolog is the yeast Δ8-Δ7 sterol isomerase, ERG2p, although the σ_1_ receptor itself has no detectable isomerase activity^11^. Human genetic data have linked point mutants in σ_1_ receptor to inherited motor neuron diseases^12–14^, and animal models implicate the receptor in Parkinson’s disease^15^, addiction^16^, and pain^17^. A σ_1_ receptor antagonist is currently in clinical trials for the treatment of neuropathic pain^7^, and agonists are in clinical trials for Alzheimer’s disease^5^ and ischemic stroke^6^.

**Fig. 1.**
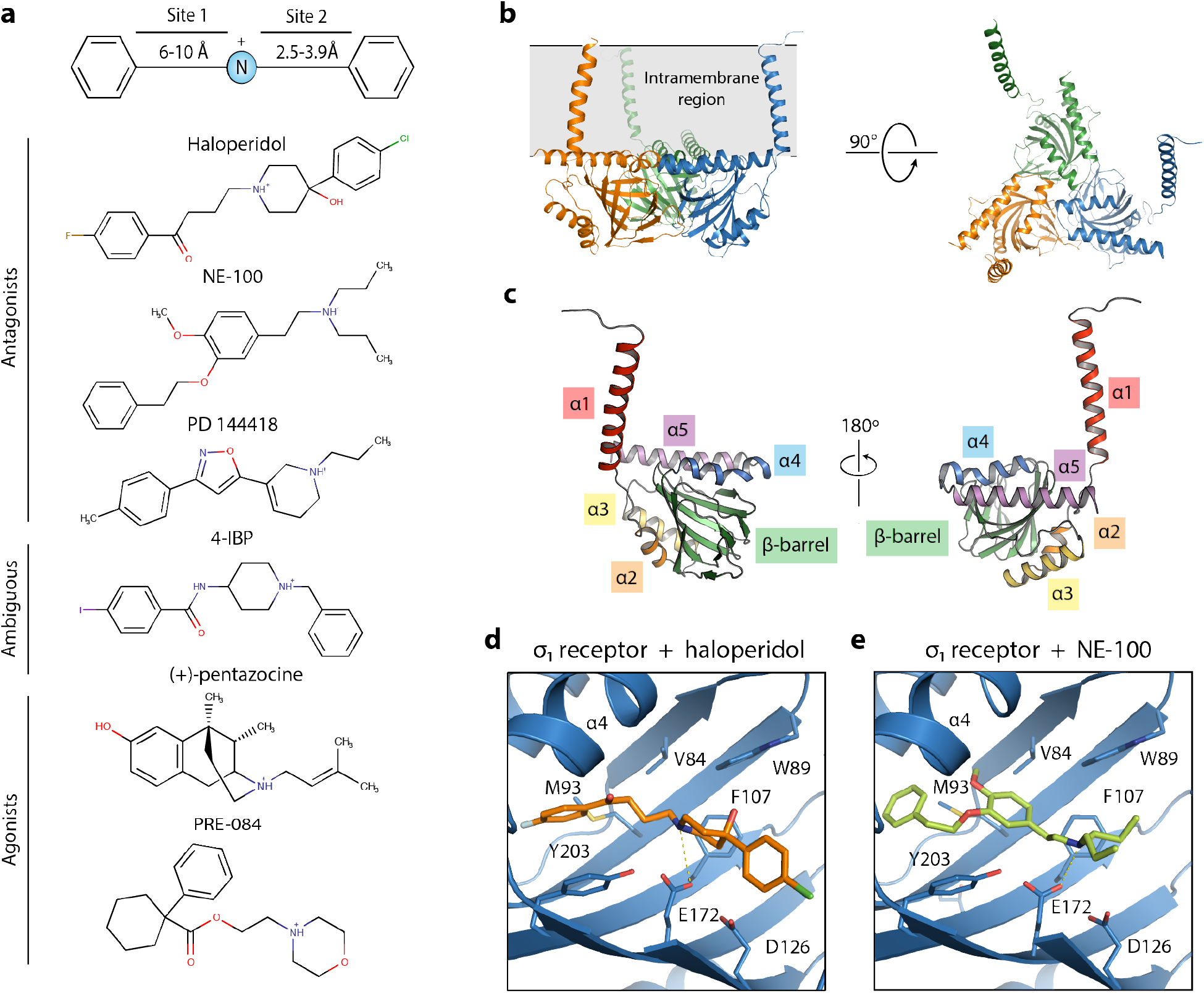
Crystal structures of human σ_1_ receptor bound to the classical antagonists haloperidol and NE-100. **a,** σ_1_ ligand pharmacophore, based on the work of Glennon *et al*.^8^ Representative σ_1_ ligands are shown below. **b,** The overall structure of the human σ_1_ receptor (PDB 5HK1). **c,** The structure of a single σ_1_ monomer, with the secondary structural elements labeled. **d** and **e,** The binding pocket of the human σ_1_ receptor (blue) binding in complex with haloperidol (orange) (**d**) and NE-100 (light green) (**e**).

Despite its potential therapeutic relevance and a wealth of high-affinity ligands, surprisingly little is known about the molecular underpinnings of σ_1_ receptor function. There is substantial evidence to suggest that the σ_1_ receptor serves as a modulator for other signaling pathway effectors^2,3,18^. Specifically, knockdown or antagonism of σ_1_ receptor can potentiate G-protein coupled receptor (GPCR) signaling^2,3^, while agonists of the σ_1_ receptor result in an IP3 receptor-dependent intracellular calcium flux^19^ and inhibition of sodium^18^ and potassium^20,21^ channel currents. The σ_1_ receptor exists in multiple oligomeric states, and reports suggest agonists causes a shift to monomeric or low molecular weight species, while antagonists bias the receptor towards high molecular weight species^4,22–24^. However, the dominant physiologically relevant oligomeric forms and the precise way in which oligomerization is tied to agonist binding are unknown.

We recently reported the first structures of the human σ_1_ receptor bound to two different ligands, PD 144418, an antagonist, and 4-IBP, a poorly characterized ligand of ambiguous efficacy class^25^. The receptor crystallized as a trimer, with each protomer showing a fold including a single transmembrane domain and a β-barrel flanked by α helices^25^ (Fig. 1b, Fig. 1c, and Supplementary Fig. 1a) While these initial results provided the first structural information on σ_1_ receptor, neither ligand is commonly used to study σ_1_ receptor function, and little functional data are available for either.

In order to understand the molecular basis for agonist activity at σ_1_, we pursued structural studies of three well-characterized classical ligands of the receptor: the antagonists haloperidol and NE-100, and the agonist (+)-pentazocine. Using the lipidic cubic phase method we determined X-ray crystal structures of the receptor in complex with these three compounds at resolutions of 3.1 Å, 2.9 Å, and 3.1 Å, respectively (Table 1, Supplementary Fig. 2).

**Table 1.** Data collection and refinement statistics.

| | $\sigma_1$ bound to<br>Haloperidol<br>(PDB 6DJZ) | $\sigma_1$ bound to<br>NE-100<br>(PDB 6DK0) | $\sigma_1$ bound to<br>(+)-pentazocine<br>(PDB 6DK1) |
| --- | --- | --- | --- |
| **Data collection** |  |  |  |
| Wavelength (Å) | 1.033 | 1.033 | 1.033 |
| Space group | P 2<sub>1</sub> 2<sub>1</sub> 2 | P 2<sub>1</sub> 2<sub>1</sub> 2 | P 2<sub>1</sub> 2<sub>1</sub> 2 |
| Number of crystals | 1 | 7 | 1 |
| Unit cell dimensions |  |  |  |
| a, b, c (Å) | 85.1, 126.1, 110.6 | 85.0, 127.0, 110.0 | 85.8, 128.6, 109.2 |
| $\alpha$, $\beta$, $\gamma$ (°) | 90, 90, 90 | 90, 90, 90 | 90, 90, 90 |
| Resolution (Å) | 50 - 3.1 (3.30 - 3.10) | 50 - 2.9 (3.00 - 2.90) | 46 - 3.1 (3.33 - 3.12) |
| Completeness (%) | 99.8 (99.6) | 95.3 (96.8) | 93.3 (82.9) |
| <I/$\sigma$(I)> | 5.9 (0.4) | 5.6 (0.5) | 3.5 (0.6) |
| CC<sub>1/2</sub> (%) | 99.4 (16.4) | 98.9 (24.0) | 99.3 (33.4) |
| Multiplicity | 5.7 (5.0) | 7.2 (7.2) | 2.5 (2.2) |
| **Refinement** |  |  |  |
| Resolution (Å) | 40.5 - 3.1 | 46.2 - 2.9 | 45.9 - 3.1 |
| No. reflections | 21454<br>(1901 in test set) | 25243<br>(2164 in test set) | 20178<br>(1673 in test set) |
| R<sub>work</sub>/R<sub>free</sub> (%) | 23.9/27.6 | 25.1/27.3 | 24.7/27.8 |
| No. atoms |  |  |  |
| Protein | 5021 | 5052 | 5054 |
| Ligand | 78 | 78 | 63 |
| Solvent ions/lipid | 164 | 155 | 204 |
| Water | 30 | 96 | 86 |
| B factors (Å<sup>2</sup>) |  |  |  |
| Protein | 102.4 | 83.0 | 78.5 |
| Ligands | 112.4 | 98.0 | 82.0 |
| Solvent ions/lipids | 127.8 | 105.3 | 102.2 |
| Water | 82.9 | 69.3 | 62.6 |
| RMS deviation |  |  |  |
| Bond length (Å) | 0.003 | 0.002 | 0.002 |
| Bond angles (°) | 0.489 | 0.463 | 0.471 |
| Ramachandran statistics |  |  |  |
| Favored | 98.3% | 98.3% | 97.7% |
| Allowed | 1.7% | 1.7% | 2.3% |
| Outliers | 0% | 0% | 0% |

## Results

### Structure of human σ_1_ receptor bound to antagonists

The structures of the σ_1_ receptor bound to the classical antagonists haloperidol and NE-100 are highly similar to each other and to our previously reported structures of σ_1_ bound to PD 144418 and 4-IBP^25^ (Supplementary Figures 1b-1e). Both haloperidol and NE-100 include a shared simple pharmacophore (Fig. 1a), and both adopt similar conformations in the ligand binding site (Fig. 1d and 1e). In each case, the ligand’s positively charged nitrogen forms an electrostatic interaction with E172, and the rest of the molecule adopts a linear pose that fits within the space not occluded by the many bulky hydrophobic residues that line the interior of the σ_1_ binding pocket (Fig. 1d and 1e). In general, the longer of the two hydrophobic regions occupies the region of the β-barrel that is proximal to the membrane, near the space between helices α4 and α5 (Fig. 1d, 1e, and Supplementary Figure 1b-e). In contrast, the shorter hydrophobic region occupies space near the bottom of the β-barrel, near D126 (Fig. 1d, 1e, and Supplementary Figure 1b-e).

### Structure of the human σ_1_ receptor bound to an agonist

The structures above reveal the overall pose of ligands in the antagonist-bound σ_1_ receptor, confirming a highly conserved binding mode and receptor conformation even for chemically diverse antagonists. Next, we investigated the structure of the receptor bound to (+)-pentazocine at 3.1 Å resolution (Fig. 2, Table 1, Supplementary Figures 2d and 2e). In general, the agonist-bound receptor crystallized similarly to antagonist-bound σ_1_, and the overall conformation of the receptor did not change significantly (Fig. 2a). The exception is a movement of helix α4, which shifts roughly 1.8 Å away from helix α5 in the (+)-pentazocine bound structure relative to the PD 144418-bound structure (Fig. 2b). This movement appears to be a consequence of the pose adopted by (+)-pentazocine, which occupies a different portion of the receptor binding pocket than the other ligands examined thus far (Fig. 2b and 2c). This difference in helix α4 position is also consistently observed in MD simulations (Fig. 2d). In simulations of unliganded σ_1_, the helix adopts a similar position as when an antagonist is bound (Fig. 2d), also suggesting that the agonist is responsible for the conformational change. (+)-pentazocine engages in an electrostatic interaction with E172, and site 2 is positioned similarly to those of the antagonists, but its nonlinear shape forces site 1 to occupy space closer to helix α4 and further from α5 relative to the antagonists. In order to prevent a steric clash between the aromatic ring of (+)-pentazocine’s benzomorphan group and residue A185 in helix α4 (Fig. 2c), helix α4 must shift towards the membrane and away from the ligand. This movement creates a slightly larger gap between helices α4 and α5 in the (+)-pentazocine-bound structure relative to the antagonist-bound structures. In the two best-resolved protomers of the (+)-pentazocine-bound structures, two water molecules occupy the space normally occupied by a portion of the antagonist.

**Fig. 2.**
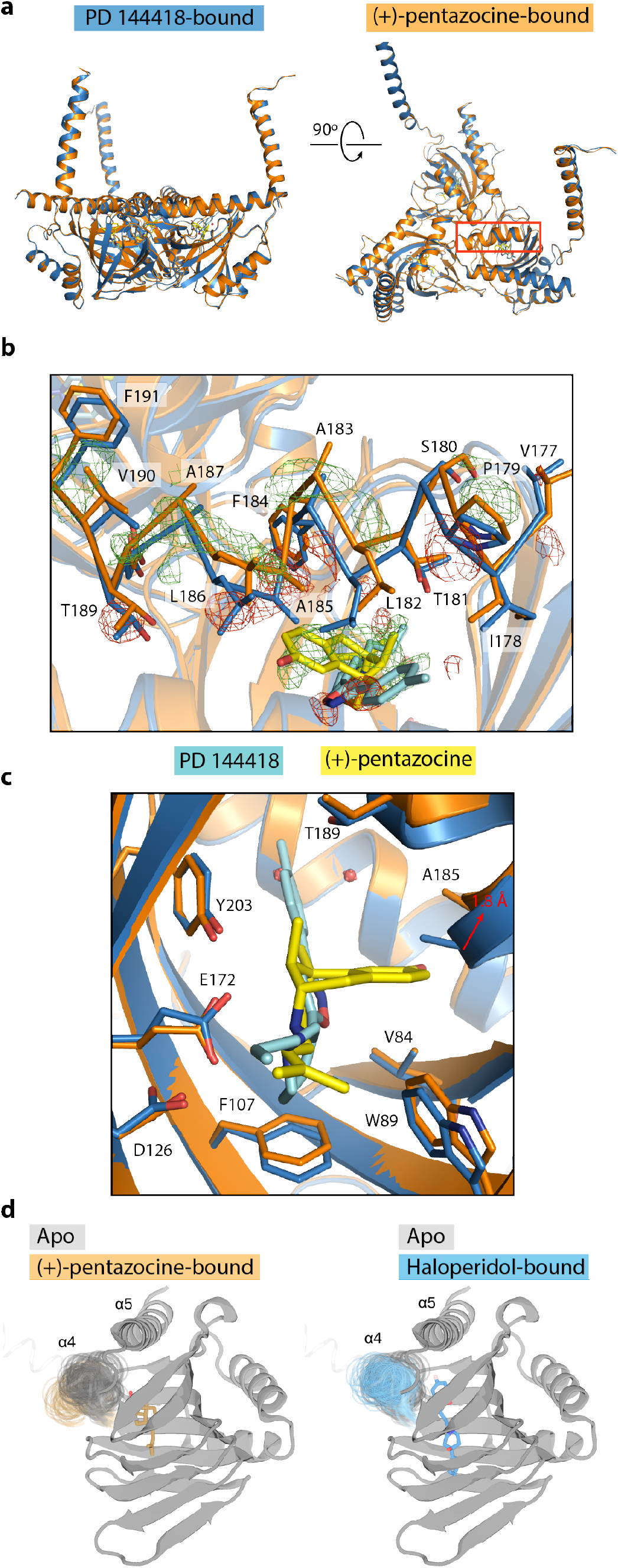
Crystal structure of the human σ_1_ receptor bound to the classical agonist (+)-pentazocine. In all panels, the structure of human σ_1_ receptor bound to (+)-pentazocine is shown in orange, and the structure of the human σ_1_ receptor bound to PD 144418 (PDB ID: 5HK1) is show in blue. The ligands (+)-pentazocine and PD 144418 are shown in yellow and cyan, respectively. Wire mesh represents Fo-Fo density, where green mesh corresponds to regions where there is more electron density in the (+)-pentazocine-bound structure relative to the PD 144418-bound structure, and red mesh is the opposite. **a,** An alignment of the overall structures of σ_1_ receptor bound to PD 144418 and (+)-pentazocine, with helix α4 highlighted by a red box. **b,** A close up of helix α4 alignment that was highlighted in a, shown in a stick representation with Fo-Fo density contoured at 2.5σ. **c,** An alignment of the (+)-pentazocine and PD 144418-bound structures in the binding pocket. **d.** Helix α4 position in simulation for unliganded (grey), (+)-pentazocine-bound (orange), and haloperidol-bound (blue) conditions. Multiple simulation frames comprising approximately 1μs of simulation per condition are shown.

### Kinetic analysis of σ_1_ receptor ligand association and dissociation

As noted above, the ligand binding site of the σ_1_ receptor is sterically occluded, and so the receptor must undergo a conformational change to allow ligand entry and egress. Previous work has shown that (+)-pentazocine associates with the receptor slowly, but rate constants for association and dissociation were not determined^26^. In order to gain a better understanding of how ligands associate and dissociate with the σ_1_ receptor, we undertook an analysis of ligand binding kinetics using [^3^H](+)-pentazocine and membranes prepared from Sf9 cells expressing σ_1_ receptor.

We began by measuring off rate at 37 °C and found it to follow a slow exponential decay with a half-life of over 200 minutes (Supplementary Fig. 3a). To obtain detailed association kinetics, we turned to a scintillation proximity assay (SPA), in which purified FLAG-tagged receptor was bound to YSi SPA beads coated with protein A and M1 αFLAG antibody. In this format, a single reaction can be monitored continuously at room temperature for an extended period (Supplementary Fig. 3b). The measured K_d_ in SPA experiments was indistinguishable from that measured in membrane binding experiments, suggesting that the receptor-ligand interaction is similar in both lipid membranes and in detergent (Supplementary Fig. 3c). These experiments showed that the association of [^3^H](+)-pentazocine to the σ_1_ receptor was not monophasic, but could be well modeled by a two-step association model, in which a zero-order reaction is followed by a concentration-dependent association step (Fig. 3a, 3b, and 3c). We also measured ligand dissociation in SPA format. In contrast to the association reaction, the dissociation data fit well to a simple monophasic dissociation curve (Fig. 3c and Supplementary Fig. 3d).

**Fig. 3.**
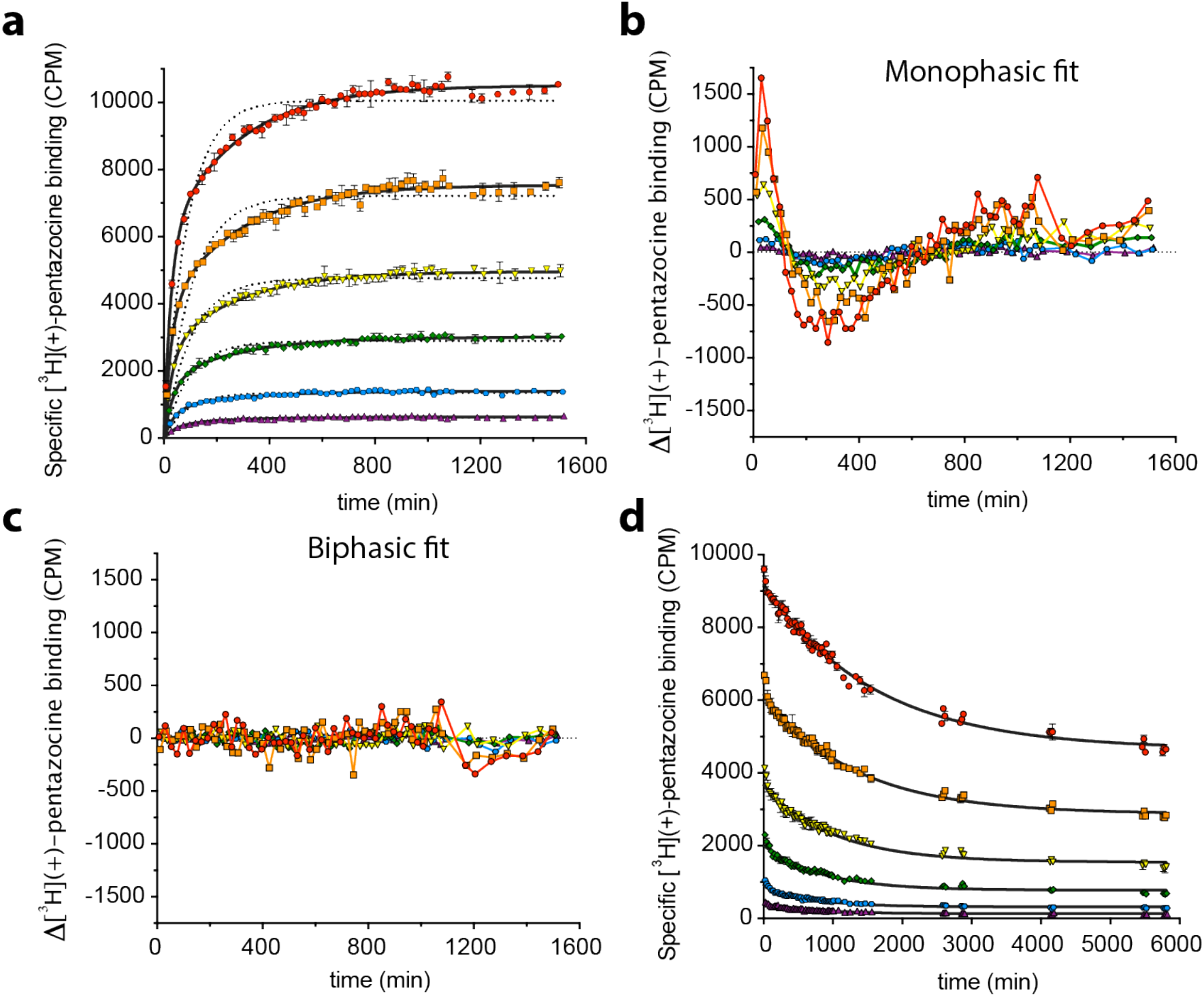
kinetic analysis of ligand binding to the σ_1_ receptor. **a,** Association of [^3^H](+)-pentazocine with the σ_1_ receptor measured in SPA format at 23 °C. The six ligand concentrations assayed are 300 nM (red), 100 nM (orange), 30 nM (yellow), 10 nM (green), 3 nM (blue), and 1 nM (violet). The best-fit monophasic curve for each concentration is shown as a dotted black line, and the best-fit biphasic curve for each concentration is shown as solid black lines. Error bars represent SEM. Data shown are representative from three independent experiments performed in duplicate. **b, c,** Residual plots for the monophasic (**b**) and biphasic (**c**) association curves and the individual data points. Colors differentiate concentration of [^3^H](+)-pentazocine used, and are the same as in **a. d,** Dissociation of [^3^H](+)-pentazocine from the σ_1_ receptor in SPA format at 23 °C. Colors represent initial [^3^H](+)-pentazocine concentrations as denoted in **a.** Solid black lines represent the best-fit monophasic exponential decay curve. Error bars represent SEM. Data shown are representative of two independent experiments performed in duplicate.

Interestingly, both the apparent k_off_ and the k_fast_ parameters for [^3^H](+)-pentazocine dissociation and association varied nonlinearly with concentration, indicative of cooperativity in ligand binding (Supplementary Fig. 3e and 3f, Table 2). However, the Hill coefficient for ligand binding in equilibrium experiments is indistinguishable from 1 (Supplementary Fig. 3g). Therefore, the binding of (+)-pentazocine to one σ_1_ monomer alters the rate of ligand binding to the next monomer, but must equally affect both on and off rates. Additionally, though the association curve for each individual concentration could be fit to a two-step exponential function, a simple two-state model is insufficient to account for the global data. This suggests that though there are at least two steps to ligand association with the σ_1_ receptor, there are probably additional steps or conformational states that are not accounted for with a simple two-step fit.

**Table 2.**
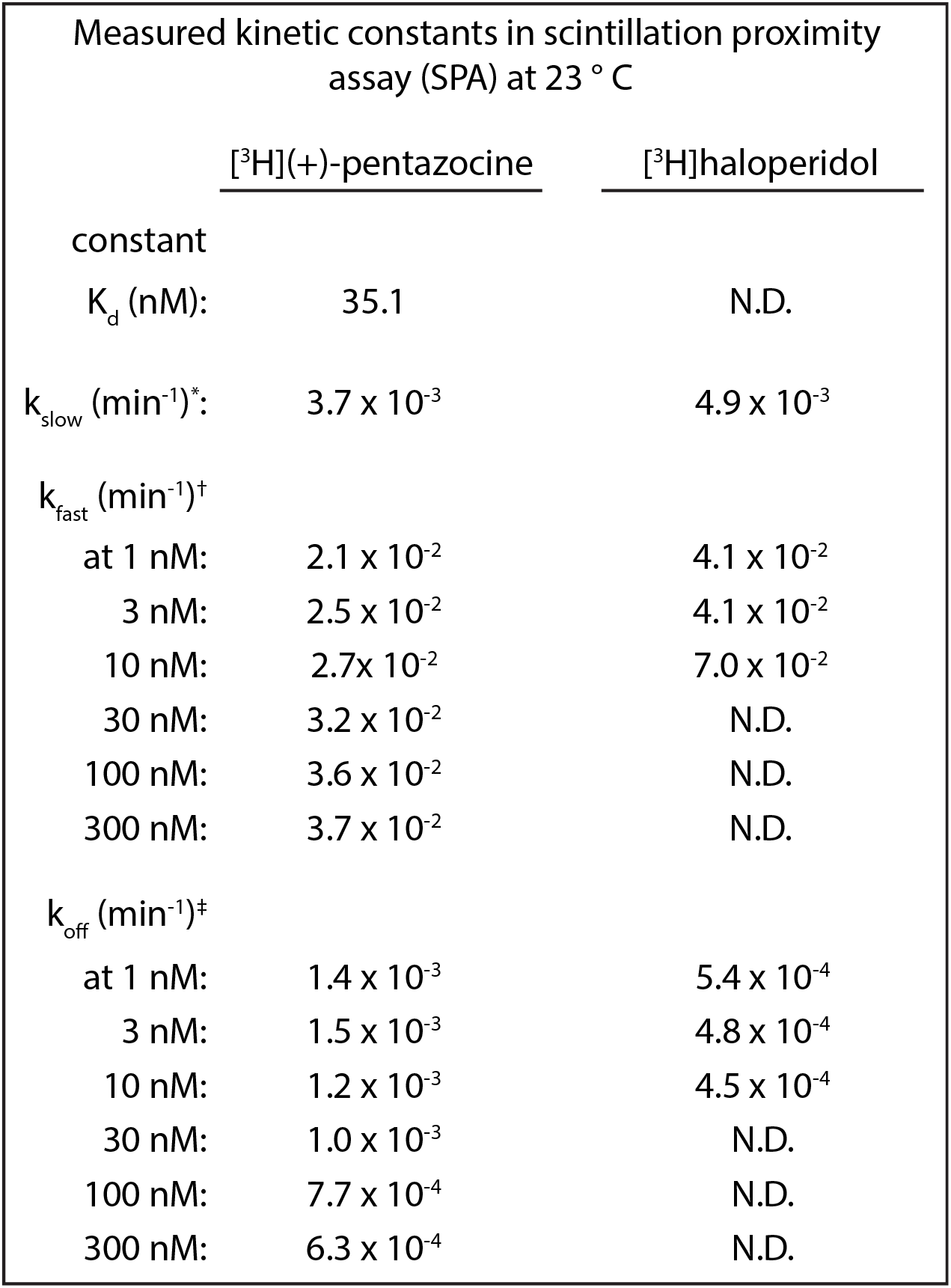
Kinetic constants for σ_1_ receptor ligand binding. ^*^k_slow_ is the observed rate constant for the slow step for ligand association to the σ_1_ receptor in the biphasic fit. ^†^k_fast_ is the observed rate constant for the fast step in ligand association to the σ_1_ receptor in the biphasic fit. ^‡^k_off_ is the observed off rate for a given concentration in the SPA measurements, based on a monophasic fit.

| Measured kinetic constants in scintillation proximity assay (SPA) at 23 ° C |  |  |
| --- | --- | --- |
|  | <u>[<sup>3</sup>H](+)-pentazocine</u> | <u>[<sup>3</sup>H]haloperidol</u> |
| constant |  |  |
| $K_d$ (nM): | 35.1 | N.D. |
| $k_{\text{slow}}$ (min<sup>-1</sup>): | $3.7 \times 10^{-3}$ | $4.9 \times 10^{-3}$ |
| $k_{\text{fast}}$ (min<sup>-1</sup>)<sup>†</sup> | | |
| at 1 nM: | $2.1 \times 10^{-2}$ | $4.1 \times 10^{-2}$ |
| 3 nM: | $2.5 \times 10^{-2}$ | $4.1 \times 10^{-2}$ |
| 10 nM: | $2.7 \times 10^{-2}$ | $7.0 \times 10^{-2}$ |
| 30 nM: | $3.2 \times 10^{-2}$ | N.D. |
| 100 nM: | $3.6 \times 10^{-2}$ | N.D. |
| 300 nM: | $3.7 \times 10^{-2}$ | N.D. |
| $k_{\text{off}}$ (min<sup>-1</sup>)<sup>‡</sup> | | |
| at 1 nM: | $1.4 \times 10^{-3}$ | $5.4 \times 10^{-4}$ |
| 3 nM: | $1.5 \times 10^{-3}$ | $4.8 \times 10^{-4}$ |
| 10 nM: | $1.2 \times 10^{-3}$ | $4.5 \times 10^{-4}$ |
| 30 nM: | $1.0 \times 10^{-3}$ | N.D. |
| 100 nM: | $7.7 \times 10^{-4}$ | N.D. |
| 300 nM: | $6.3 \times 10^{-4}$ | N.D. |

Since the rate-limiting step for [^3^H](+)-pentazocine association was not dependent on ligand concentration, we suspected that this step represented a conformational change from a ligand-inaccessible to a ligand-accessible state. To test this hypothesis, we repeated the experiment with [^3^H]haloperidol. The association of [^3^H]haloperidol to the σ_1_ receptor was also poorly modeled by a one-step reaction, but fit well to a two-step model (Supplementary Fig. 3h and 3i). Additionally, the rate of the slow step was essentially identical for both [^3^H]haloperidol and [^3^H](+)-pentazocine (Table 2), which is consistent with the conclusion that this step represents a conformational change intrinsic to the receptor that is ligand-independent. As seen with [^3^H](+)-pentazocine, dissociation of [^3^H]haloperidol from σ_1_ receptor was slow and could be modeled by a monophasic exponential decay function (Supplementary Fig. 3j and 3k).

### Ligand binding pathway via molecular dynamics simulation

To better characterize the pathway of ligand binding and dissociation, we performed MD simulations of σ_1_ with the goal of characterizing possible conformational rearrangements that could expose the binding pocket. To reduce the computational complexity of the system, we simulated the σ_1_ monomer and used accelerated molecular dynamics, which applies a boost to dihedral energy minima in order to speed up observation of conformational changes^27^.

The σ_1_ monomer from the crystal structure was inserted into a hydrated lipid bilayer, with (+)-pentazocine removed from the binding pocket and placed in the water. Using multiple rounds of simulation totaling over 110 μs, we were able to assemble a three-step binding pathway, with (+)-pentazocine reaching a bound state with an RMSD < 3 Å to the crystallographic pose (Fig. 4a).

**Fig. 4.**
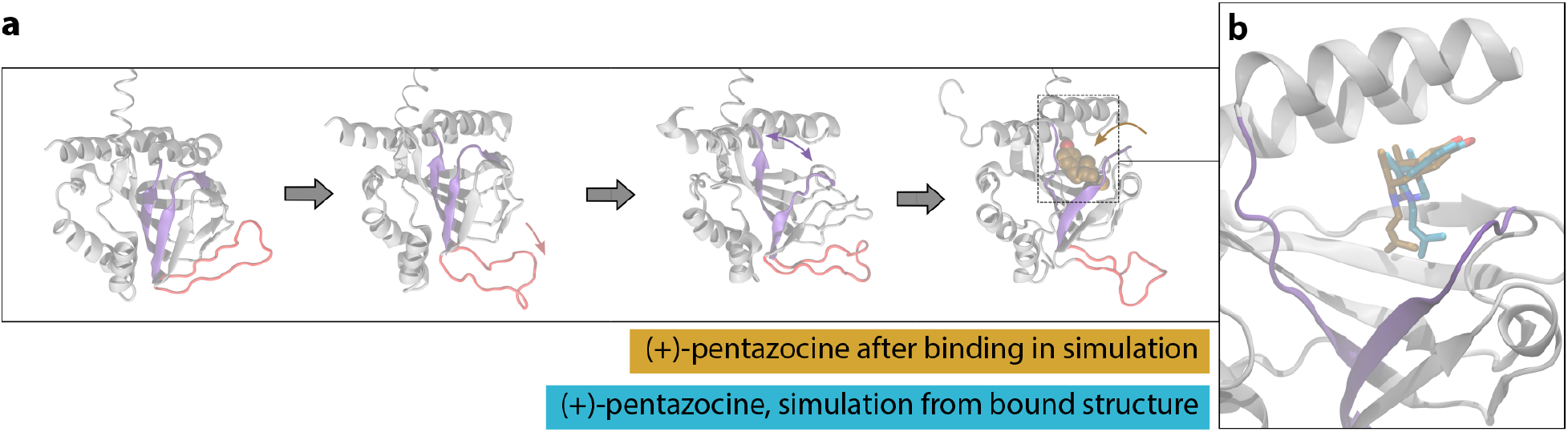
Molecular dynamics simulation reveals a putative binding pathway for (+)-pentazocine. **a.** Binding pathway of (+)-pentazocine, with the “lid” region shown in red and the beta strands that separate shown in purple. From left to right: simulation begins with an unliganded receptor, where the “lid” region then opens. Next, the interior of the receptor opens, and the ligand enters and binds through this opening. **b.** A simulation frame after the ligand has entered the binding pocket (tan), compared to a frame initiated from the crystal structure with the ligand bound (blue). Protein backbone is shown for the binding simulation in grey. Helix α4 is located at the top of the rendering.

This binding pathway requires two major conformational rearrangements in order for the pocket to become accessible to the ligand. First, the “lid” of the receptor opens, breaking backbone hydrogen bonds between Trp136 and Ala161. Next, the beta-barrel structure in the interior of the receptor separates, breaking backbone hydrogen bonds between Glu123 and Arg175 and exposing the binding pocket. The ligand enters through this opening near the membrane, and assumes a near-crystallographic pose as the protein closes around it (Fig. 4b).

Each of these rare conformational changes or binding events was observed multiple times in simulation. Interestingly, the beta-barrel separation that exposes the binding site was only observed in simulations where the receptor “lid” had already opened, suggesting that two sequential conformational changes may be necessary before the ligand can bind. The lid opening may be a prerequisite for further conformational change as it may perturb the internal hydrogen-bond network of σ_1_ so that larger rearrangements may occur.

## Discussion

Antagonism or genetic ablation of σ_1_ receptor has analgesic effects in whole animals and humans^7,17,28^, and potentiates GPCR signaling in cells^2,3^. In contrast, σ_1_ receptor agonists are usually defined by their ability to oppose the effects of agonists, and have been associated with cytoprotective effects^29–31^. Currently, the biochemical basis for agonism or antagonism at the σ_1_ receptor is largely unknown, which complicates the unambiguous assignment of efficacy class for σ_1_ ligands. The most well-documented biochemical difference between the two ligand classes is that antagonists increase the receptor’s oligomeric state, while agonists decrease the oligomeric state^4,22–24^. The structural data we present here show that these ligands occupy a different region of the binding pocket (Fig. 2c and 2d). Antagonists adopt a more linear pose, with the primary hydrophobic region of the molecule pointing towards the space between helices α4 and α5, while (+)-pentazocine’s primary hydrophobic site points towards helix α4 (figures 2c and 2d). Presumably, structurally similar agonists like (+)-SKF-10,047 adopt a similar pose, accounting for their shared biological activities.

As a result of the steric constraints of agonists, most of helix α4 is forced to shift 1.1-1.8 Å away from helix α5 to accommodate the ligand. In our structure, this shift in α4 does not disrupt the oligomerization interface between individual protomers. However, if α4 were to move to a greater degree, it could disrupt the oligomerization interface. This is consistent with prior data, which suggest that σ_1_ receptor agonists bias the receptor towards lower molecular weight states, while antagonists bias it towards higher molecular weight states^4, 22–24^. Additionally, molecular modeling by Yano *et al*. predicted that (+)-pentazocine and other multimer-impeding ligands would occupy this space differentially from haloperidol and other multimer-promoting ligands^24^, which is consistent with our structural results. Importantly, crystallographic studies by necessity favor conformationally stable, low-energy states, and so the structures shown here may not represent a fully activated state of the receptor. Indeed, studies of G protein-coupled receptors bound to agonists often show inactive-state structures in the absence of G proteins or antibody fragment stabilizers^32^.

We have also shown by kinetic analysis that (+)-pentazocine associates with the σ_1_ receptor in at least two steps, with MD simulation suggesting a three-step process requiring two substantial conformational changes to the receptor. Ligand association and dissociation at the σ_1_ receptor is very slow, and the rate-limiting step is independent of ligand concentration. Though many groups have analyzed the effects of σ_1_ receptor ligands in cells, there is no standard incubation time for observing σ_1_-dependent effects of σ ligands. A brief survey of the literature reveals that when using σ_1_ receptor ligands in cellular or biochemical assays, incubation times and temperatures vary from room temperature for 20 minutes^33^ to 37 °C incubation for up to 72 hours^31^. Ligand concentrations are sometimes nearly 10,000-fold over Kd^34–36^. Our data indicate that it can take 1.5 hours or longer to reach saturation at 37 °C, and at room temperature it can take nearly a day. Furthermore, since the rate-limiting step is concentration-independent, high ligand concentrations cannot overcome the receptor’s slow binding kinetics. Therefore, when ascribing the effects of σ_1_ receptor ligands to the σ_1_ receptor, one must ensure that sufficient time is allowed for the ligands to engage with the receptor. If effects are observed too quickly and are only observed at ligand concentrations that are vastly higher than Kd, then it is unlikely that the effects are σ_1_-mediated.

We have shown that the agonist (+)-pentazocine adopts a binding pose in the σ_1_ receptor binding pocket that is different from that of antagonists, which tend to bind similarly to one another despite their chemical diversity. We have also demonstrated that ligands associate with the σ_1_ receptor very slowly and in multiple steps. Our simulations suggest that ligands enter the binding pocket through a dynamic opening that would be challenging to predict based on a single crystal structure.

However, the precise details of σ_1_ signaling in cells have yet to be determined. While there are myriad proposed binding partners for the σ_1_ receptor^1,3,4,19,30^, the critical effectors of σ_1_ receptor signaling still need to be unambiguously established. Future work will need to focus on these functional questions in order to fully understand the function of the σ_1_ receptor and its potential as a therapeutic target.

## Online Methods

### Protein expression and purification

Human σ_1_ receptor and was expressed and purified in Sf9 cells in a manner similar to that described previously^25^. In brief, the receptor was cloned into pFastBac1 with an amino-terminal hemagglutinin signal sequence, followed by a FLAG epitope tag and a 3C protease cleavage site. The receptor was expressed in Sf9 cells (Expression Systems) using the FastBac baculovirus system (ThermoFisher). Cells were grown in a shaker at 27 °C and infected when they had reached a density of 4 x 10^6^ cells/mL. After infection, the cells were allowed to grow for 48-52 h, at which point they were harvested for centrifugation. Pellets were stored at −80 °C until use.

For samples used in crystallography, 1 μM of either haloperidol, (+)-pentazocine, or NE-100 was added to all purification steps. Haloperidol and NE-100 were purchased from Tocris Biosciences, and (+)-pentazocine was kindly provided by Dr. Felix Kim. For samples used for SPA binding assays, no ligand was added. Cell pellets were thawed and lysed by osmotic shock in 20 mM HEPES pH 7.5, 2 mM MgCl2, and 1:200,000 (v/v) benzonase nuclease (Sigma Aldrich). The lysate was then spun at 48,000 x g for 20 min at 4 °C. The supernatant was discarded, and the pellets were solubilized using a glass dounce tissue homogenizer in a buffer containing 1% (w/v) lauryl maltose neopentyl glycol (LMNG, Anatrace), 20 mM HEPES pH 7.5, 250 mM NaCl, and 20% glycerol. For samples used in the crystallization of σ_1_ receptor with haloperidol and (+)-pentazocine, the solubilization buffer also contained 0.1% (w/v) cholesterol hemisuccinate (CHS; Stealoids). However, samples used in the crystallization of σ_1_ receptor with NE-100 and in SPA experiments were not solubilized with CHS, as it was found to have no effect on protein quality or yield. Following homogenization, samples were stirred at 4 °C for 2 h, and then centrifuged again as before. The supernatant was filtered over glass microfiber filters, and 2 mM CaCl_2_ was added to the solution. The sample was then run over 4 mL of αFLAG affinity resin. Once the sample was loaded on the resin, it was washed once with 50 mL of buffer containing 0.1% LMNG, 20 mM HEPES pH 7.5, 150 mM NaCl, 2% glycerol, and 2 mM CaCl_2_, then again with a buffer containing 0.01% LMNG, 20 mM HEPES pH 7.5, 150 mM NaCl, 0.2% glycerol, and 2 mM CaCl_2_. Following these wash steps, the protein was eluted in an identical buffer that lacked CaCl_2_ but was supplemented with 0.2 mg/mL FLAG peptide and 5 mM EDTA. Samples used for crystallography were incubated with 3C protease at 4 °C overnight to remove the FLAG tag. Samples used for SPA experiments were also left at 4 °C overnight but were not digested.

The next day, samples were further purified by SEC on a Sephadex S200 column (GE Healthcare). The buffer for SEC contained 0.01% LMNG, 20 mM HEPES pH 7.5, and 150 mM NaCl. For samples used for crystallography, the buffer also contained 1 μM of the desired ligand. After SEC, samples intended for crystallization were concentrated to 25-35 mg/mL and flash frozen in liquid nitrogen in 8-9 μL aliquots. Samples intended for SPA experiments were concentrated to 300-400 μM and diluted to 200 μM in SEC buffer supplemented with 20% glycerol. The SPA samples were aliquoted into 6 μL aliquots and flash frozen in liquid nitrogen. All samples were stored at −80 °C and never frozen again after thawing.

### Crystallography and data collection

Purified σ_1_ receptor was reconstituted into lipidic cubic phase as described previously^25,37^. The cubic phase was dispensed in 30 nL drops onto a hanging drop cover and overlaid with 600 nL of precipitant solution with the use of a Gryphon LCP robot (Art Robbins Instruments). The crystal that provided the haloperidol-bound structure was grown in 500 mM Li_2_SO_4_, 35% (v/v) PEG 300, 1% (v/v) hexanediol, and 100 mM MES pH 6.4. The crystal that provided the (+)-pentazocine-bound structure was grown in 240 mM Li_2_SO_4_, 42% (v/v) PEG 300, 1% (v/v) hexanediol, and 0.1 M MES pH 6.0. The crystals for the NE-100 bound dataset were grown in 400-500 mM Li_2_SO_4_, 30-40% PEG 300, 1% hexanediol, and 0.1 M MES pH 5.8-6.0. Crystals grew slowly over the course of one to three weeks, and were harvested with mesh loops (MiTeGen) and stored at −80 °C. For (+)-pentazocine-bound crystals, it was important to harvest within two weeks, or crystal quality would decline substantially.

Data collection was performed at Advanced Photon Source GM/CA beamlines 23ID-B (NE-100 and (+)-pentazocine-bound structures) and 23ID-D (haloperidol-bound structure). Data collection was performed as described previously^25^. The datasets for haloperidol-bound and (+)-pentazocine-bound σ_1_ receptor were obtained from single crystals, while the dataset for NE-100-bound σ_1_ receptor was obtained by merging partial datasets from seven crystals.

### Data processing, structure refinement, and model building

Data were processed using XDS^38^. For the haloperidol and NE-100 bound complexes, scaling was done with XSCALE^38^. For the (+)-pentazocine-bound structure, scaling was done with Aimless^39^. Phases for all three structures were solved via molecular replacement, using PD144418-bound σ_1_ receptor (PDB ID: 5HK1) as a search model. Model building was done with Coot^40^, and refinement was performed in phenix.refine^41^. Following refinement, structures were evaluated with MolProbity^42^, and figures were prepared with PyMOL^43^. The SBGrid Consortium supported all crystallographic data processing, refinement, and analysis software^44^.

### Preparation of membranes for radioligand binding

Membranes were prepared as described previously^45^, using a protocol adapted from that of Vilner *et al*.^46^. In brief, Sf9 cells expressing σ_1_ receptor were harvested by centrifugation and lysed by osmotic shock in a buffer containing 20 mM HEPES pH 7.5, 2 mM MgCl2,1:200,000 (v/v) benzonase nuclease (Sigma Aldrich), and cOmplete Mini EDTA-free protease inhibitor tablets (Sigma Aldrich). The lysates were homogenized using a glass dounce tissue homogenizer and then centrifuged at 48,000 x g for 20 min. Following centrifugation, the membranes were resuspended in buffer containing 50 mM Tris pH 8.0 and cOmplete Mini EDTA-free protease inhibitor tablets (Sigma Aldrich). The samples were spun down as before and resuspended in the same buffer. Next, the samples were homogenized using a needle and syringe. Protein content was determined using the DC protein assay (Bio Rad). Samples were aliquoted into 100 μL aliquots at protein concentrations of 10-20 mg/mL and flash frozen in liquid nitrogen. All samples were stored at −80 °C until use.

### Saturation binding in Sf9 membranes

Saturation binding was performed as described previously^45^, using a method similar to that of Chu and Ruoho^47^. Briefly, membrane samples from Sf9 cells expressing wild-type or mutant σ_1_ receptor prepared as described above were thawed, homogenized with a syringe, and diluted in 50 mM Tris pH 8.0. Each reaction was 100 μL, with a final concentration of 0.025 mg/mL protein and the indicated concentration of [^3^H](+)-pentazocine. To assay nonspecific binding, equivalent reactions containing 2 μM haloperidol were performed in parallel. Samples were shaken at 37 °C for 90 minutes. Afterwards, the reaction was terminated by massive dilution and filtration over a glass microfiber filter using a Brandel harvester. Filters were soaked with 0.3% polyethyleneimine (PEI) for at least 30 minutes before use. Radioactivity was measured by liquid scintillation counting.

### Measurement of ligand dissociation in Sf9 membranes

Membrane samples prepared as described above were thawed, syringe homogenized, and diluted in 50 mM Tris pH 8.0 to a final concentration of 0.05 mg/mL in a 96-well plate with a final volume of 100 μL per well. To equilibrate the samples with radioligand, samples were incubated with 10 nM [^3^H](+)-pentazocine (Perkin Elmer) for 90 minutes at 37 °C. After equilibration, 1 μL 500 μM haloperidol (Tocris Biosciences) was added to a set of wells in triplicate for a particular time point. This was repeated for a total of eight time points over the course of twenty-four hours. After twenty-four hours, the reaction was terminated by massive dilution in ice-cold water and filtration over a glass microfiber filter using a Brandel harvester. Filters were soaked in 0.3% PEI for at least 30 minutes prior to use. Radioactivity was quantified by liquid scintillation counting. Data analysis was performed using GraphPad Prism.

### Scintillation proximity assay

All scintillation proximity experiments were performed using protein-A coated YSi scintillation proximity beads (PerkinElmer, RPN143). Beads coupled with M1 αFLAG antibody and stored in HBS at 4 °C until use in 5 mg aliquots. Upon use, 4-6 mg of beads were spun down twice in a cold centrifuge and resuspended each time in HBS with 0.01% LMNG and 2 mM CaCl_2_. Then, the beads were incubated with 50 nM σ_1_ receptor purified as described above for 30 min at 4 °C. Following coupling of the receptor, the beads were again centrifuged and resuspended twice in HBS with 0.01% LMNG and 2 mM CaCl_2_. To start the reaction, 40 μL containing 0.2 mg of receptor-linked beads was added to a solution containing the desired concentration of radioligand in a total volume of 360 μL, for a total reaction volume of 400 μL. To assay nonspecific binding, equivalent reactions were prepared that also contained either nonradioactive haloperidol ([^3^H](+)-pentazocine binding) or nonradioactive NE-100 ([^3^H]haloperidol binding) at a concentration of 5 μM. Once association measurements were completed, 5 μM haloperidol ([^3^H](+)-pentazocine binding) or NE-100 ([^3^H]haloperidol binding) was added to each vial to begin the dissociation measurements. Samples were measured in duplicate at room temperature using a Beckman Coulter LS 6500 multi-purpose scintillation counter. In order to average duplicate points, both the signal in CPM and the time at which the two different vials were measured were averaged. Data were analyzed using GraphPad Prism.

### Molecular dynamics simulations

#### MD simulations setup

Simulations of the σ_1_ receptor were based on either the (+)-pentazocine or haloperidol-bound crystal structures described in this manuscript. The receptor was simulated in four distinct conditions (Supplementary Table 1): (A) the (+)-pentazocine bound structure, (B) the haloperidol bound structure, (C) the (+)-pentazocine bound structure with ligand removed, and (D) the (+)-pentazocine bound structure with ligand removed and placed in solvent. All simulations were of a σ_1_ monomer.

Coordinates were prepared by first removing chains B and C to obtain a monomer, and removing crystallographic ligands. For conditions C and D, the ligand was also removed from the binding pocket and for condition D it was replaced at least 10 Å from the protein. Prime (Schrodinger, Inc) was used to model in missing side chains, add hydrogens, and cap the protein chain termini with the neutral groups acetyl and methylamide.

As suggested previously^25^, Glu172 should be charged to interact with the ligand, and Asp126 should be protonated to hydrogen bond with Glu172; the protonation states of these two residues were set accordingly. All other residues were left in their dominant protonation state at pH 7.0.

The prepared protein was aligned to the Orientation of Proteins in Membranes (OPM) structure of PDB 5HK1, and internal waters added with Dowser. The structures were then inserted into a pre-equilibrated palmitoyl-oleoyl-phosphatidylcholine (POPC) bilayer, and solvated with 0.15 M NaCl in explicitly represented water, then neutralized by removing sodium ions using the program Dabble. Final system dimensions were about 63 × 63 × 140 Å^3^, including about 140 lipids, 12000 water molecules, 34 sodium ions, and 27 chloride ions.

#### MD simulation force field parameters

We used the CHARMM36m parameter set for protein molecules, the CHARMM36 parameter set for lipid molecules and salt ions, and the CHARMM TIP3P water model. Parameters for (+)-pentazocine and haloperidol were generated using the CHARMM General Force Field (CGenFF) with the ParamChem server, version 1.0.0. Full parameter sets are available upon request.

#### MD simulation protocol

Simulations were performed on GPUs using the CUDA version of PMEMD (Particle Mesh Ewald Molecular Dynamics) on Amber16. Prepared systems were minimized, then equilibrated as follows: the system was heated using the Langevin thermostat from 0 to 100 K in the NVT ensemble over 12.5 ps with harmonic restraints of 10.0 kcal·mol^−1^ Å^−2^ on the non-hydrogen atoms of lipid, protein and ligand, with initial velocities sampled from the Boltzmann distribution. The system was then heated to 310 K over 125 ps in the NPT ensemble with semi-isotropic pressure coupling and a pressure of 1 bar. Further equilibration was performed at 310 K with harmonic restraints on the protein and ligand starting at 5.0 kcal·mol^−1^ Å^−2^ and reduced by 1.0 kcal·mol^−1^ Å^−2^ in a stepwise fashion every 2 ns, for a total of 10 ns of additional restrained equilibration.

Five independent simulations were initialized from the final snapshot of the restrained equilibration for each condition (Supplementary Table 1). These simulations were conducted in the NPT ensemble at 310 K and 1 bar, using a Langevin thermostat and Monte Carlo barostat. In each of these simulations we performed 5 ns of unrestrained equilibration followed by 0.8 - 6.7 μs production run. Simulations used periodic boundary conditions and a time step of 4.0 fs with hydrogen mass repartitioning. Bond lengths to hydrogen atoms were constrained using SHAKE. Nonbonded interactions were cut off at 9.0 Å, and long-range electrostatic interactions were computed using the particle mesh Ewald (PME) method with an Ewald coefficient β of approximately 0.31 Å and B-spline interpolation of order 4. The FFT grid size was chosen such that the width of a grid cell was approximately 1 Å. Accelerated molecular dynamics (aMD) was used to boost dihedral potential energies, with parameters E_D_=10427 and α_D_=170.

For condition C, the ensemble of simulations was periodically visualized for major conformational change in the protein, and new sets of 5–10 simulation replicates initialized from restart files corresponding to rare events, with velocities either retained or equilibration being performed once more (Supplementary Table 1)

#### MD simulation analysis protocols

Trajectory snapshots were saved every 200 ps during production simulations. Trajectory analysis was performed using VMD and CPPTRAJ, and visualization performed using VMD.

## Acknowledgments

We thank B. Kelly for performing preliminary simulations of the σ_1_ receptor trimer, and Advanced Photon Source GM/CA beamline staff for excellent technical support of crystallographic data collection. We also thank Dr. Felix Kim for generously providing (+)-pentazocine for crystallographic studies. This work was supported by a Klingenstein-Simons Fellowship in Neuroscience (A.C.K.), National Institutes of Health grant 1R01GM119185 (A.C.K.), the Winthrop Fund/Harvard Brain Science Initiative (A.C.K.) and National Science Foundation Graduate Research Fellowship award number DGE1745303 (H.R.S.).

